# A robust core architecture of functional brain networks supports topological resilience and cognitive performance in aging

**DOI:** 10.1101/2022.02.07.479418

**Authors:** William Stanford, Peter J. Mucha, Eran Dayan

## Abstract

Aging is associated with gradual changes in cognition, yet some individuals exhibit protection against aging-related cognitive decline. The topological characteristics of brain networks that support protection against cognitive decline in aging are unknown. Here, we investigated whether the robustness of brain networks, queried via the delineation of the brain’s core network structure, supports superior cognitive performance in healthy aging individuals (n=320, ages 60-90). First, we decomposed each subject’s functional brain networks using k-shell decomposition, finding that cognitive function is associated with more robust connectivity of core nodes, primarily within the frontoparietal control network (FPCN). Next, we find that the resilience of core brain network nodes, within the FPCN in particular, relates to cognition. Finally, we show that the degree of segregation in functional networks mediates relationships between network resilience and cognition. Together, these findings suggest that brain networks balance between robust core connectivity and segregation to facilitate high cognitive performance in aging.

## 1. Introduction

The populations of countries all across the globe are aging (Economic and Social Affairs 2020). As populations age, dementia and other neurological conditions associated with cognitive decline are expected to become more common (Sleeman, de Brito et al. 2019). However, cognitive decline also occurs in healthy aging with significant variance between individuals (Novotný, Gonzalez-Rivas et al. 2021). Namely, some individuals are able to resist the effects of age to maintain high cognitive function late in life (Borelli, Carmona et al. 2018). With aging populations that are expected to remain in the workforce longer than previous generations, it is important to identify mechanisms responsible for supporting high cognitive performance late in life.

One common approach for discovering the mechanisms that support cognitive performance in aging is by studying brain network connectivity with tools developed in network science (Rubinov and Sporns 2010). Studies thus far have typically focused on measures of network topology, identifying age-associated changes throughout normal (Fair, Dosenbach et al. 2007, Betzel, Byrge et al. 2014, Chan, Park et al. 2014, Sala-Llonch, Bartrés-Faz et al. 2015, Cohen and D’Esposito 2016, Sadiq, Langella et al. 2021) and pathological aging (Chen, Necus et al. 2021, Langella, Sadiq et al. 2021). However, understanding how measures of network topology relate to the resilience of brain networks in aging is difficult due to lack of suitable longitudinal datasets and the inability to perform lasting experimental perturbations in human subjects. To overcome this issue, one can simulate network perturbations and quantify the network’s resilience via functional or topological measures (Albert, Jeong et al. 2000). In the context of brain networks, this is commonly done by simulating the removal of regions with high network centrality and quantifying the network’s ability to remain connected despite these perturbations (Albert, Jeong et al. 2000, Achard, Salvador et al. 2006, Bullmore and Sporns 2009, Piraveenan, Thedchanamoorthy et al. 2013). Studies that utilize these so called targeted attacks can simulate the types of pathological lesions that commonly occur in neurodegenerative diseases (Crossley, Mechelli et al. 2014), and have revealed differences in network resilience between healthy and diseased states (He, Chen et al. 2008, Mancini, De Reus et al. 2016, Cascone, Langella et al. 2021).

In quantifying network resilience to targeted attacks, measuring if the network remains connected may fall short in that the requirement of “connectedness” gives no additional information on the robustness of the underlying network connectivity before or after attacks have been performed (Mohseni-Kabir, Pant et al. 2021). An alternative approach could be to study the resilience of network features revealed by a technique known as *k*-shell decomposition (Pittel, Spencer et al. 1996, Dorogovtsev, Goltsev et al. 2006, Dorogovtsev, Goltsev et al. 2008, Kitsak, Gallos et al. 2010, Min, Morone et al. 2016). In this approach, networks are decomposed into shells of nodes, where each successive shell requires more robust interconnectivity than the previous, until a core set of nodes within a network is reached (Pittel, Spencer et al. 1996, Dorogovtsev, Goltsev et al. 2006). Next, attacks are performed on these networks, and the resilience of nodal shell and core assignments are measured by reapplication of *k*-shell decomposition (Goltsev, Dorogovtsev et al. 2006, Lee, Jo et al. 2016, Yuan, Dai et al. 2016, Schmidt, Pfister et al. 2019, Shang 2019, Mohseni-Kabir, Pant et al. 2021, Wang, Li et al. 2021, Zhou, Lv et al. 2021). Performing *k*-shell decomposition in tandem with attack simulations has yielded insights into robustness within ecological and financial networks (Burleson-Lesser, Morone et al. 2020), and facilitated the understanding of stability in social networks (Zhang, Zhang et al. 2017). Additionally, *k-*shell decomposition has been used to identify hierarchical structure, and the existence of core networks within the healthy adult brain (Hagmann, Cammoun et al. 2008, Lahav, Ksherim et al. 2016, Lucini, Del Ferraro et al. 2019). However, this approach has not yet been used to identify brain network robustness in cognitive aging, nor has it been applied in tandem with attack simulations to study resilience of brain networks.

In this study, we aimed to uncover the relationships between network resilience and healthy aging. We used data from subjects between the ages of 60-90 years old within the HCP-Aging dataset (Harms, Somerville et al. 2018) to construct functional brain networks according to the Schaefer Local-Global parcellation (Schaefer, Kong et al. 2018) from measures of resting-state functional connectivity (RSFC) (**Figure 1A**). We binarized networks using a method that selects Orthogonal Minimum Spanning Trees (OMSTs), similar to the one proposed in (Dimitriadis, Antonakakis et al. 2017) and then thresholded by edge frequency (**Figure 1B**), see **methods** for more details. Next, we applied *k*-shell decomposition to the resulting brain networks and studied the associations between robustly connected network nodes and cognition. We focused on episodic memory, a cognitive measure that is widely impacted by aging (Leal and Yassa 2015). We examined the specificity of the results by testing associations with a non-memory cognitive function (processing speed). We then simulate targeted attacks, and calculate the *k*-shell decomposition after every attack (**Figure 1C**). We hypothesized that this method of measuring resilience would show high sensitivity to differences in network topology of brain networks that confer network resilience and relate to episodic memory in healthy aging. Finally, we use system segregation (Chan, Park et al. 2014, Wig 2017, Chan, Han et al. 2021), a topological feature known to be altered in aging, to study relationships between resilience of core network nodes, the global topological organization of individual brains, and episodic memory (**Figure 1D**).

**Figure 1.**
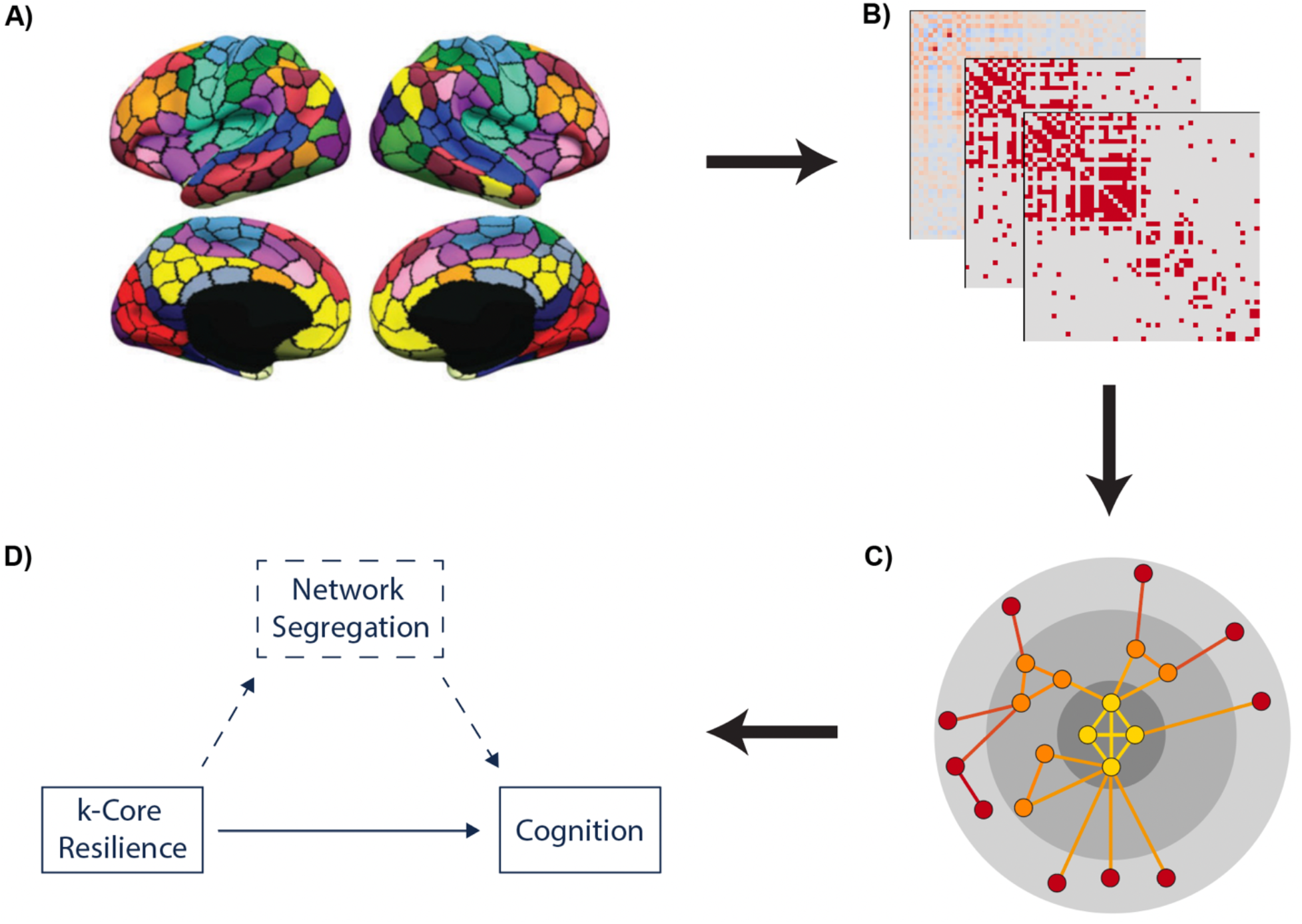
Study outline. **A)** functional MRI data from 320 subjects from the HCP aging dataset, ages of 60-90, were used for our study. Resting state functional connectivity data was registered to the Schaefer Local-Global parcellation with 17 networks and 400 ROIs. **B)** Fisher z-transformed functional connectivity matrices were thresholded using a 2-step procedure. First 20 orthogonal minimum spanning trees (OMSTs) were generated on each subject’s brain network. Second, edges were kept if they occurred greater than 50^th^ quantile of frequencies among all subjects (see **methods** for more details). **C)** We decomposed the brain networks using *k*-shell decomposition, and then quantified the resilience of each subject’s brain network using attack simulations. **D)** Finally, we investigated how robustly connected nodal cores support high cognitive performance, and the relationships between resilience of nodal cores, network segregation, and cognition.

## 2. Results

### 2.1 k-shell decomposition of brain networks

We first applied *k*-shell decomposition to each subject’s respective functional brain network. In *k*-shell decomposition, each node within a network is assigned to a specific *k*-shell (**Figure 2A**). This *k* value is determined by the maximum value *k* at which a node remains in the network while iteratively removing all nodes with degree less than *k*. (**Figure 2B**). A node in shell *k* is also in core *k*. However, a *k-*core contains all nodes in *j*-shells where *j* ≥*k* (**Figure 2C**). For more information on the calculation of a network’s *k*-shell decomposition, see **methods**. *k*-shell decomposition reveals hierarchical groups of nodes within functional brain networks (Lahav, Ksherim et al. 2016), arranged according to patterns of generally increasing connectivity. We began the analysis by calculating the median shell assignment for each region of interest (ROI) in the network across subjects (**Figure 3A**). We observed that lower shells were composed of primarily limbic ROIs (**Figure 3C**), mid-ranged shells consisted of mostly somatomotor ROIs (**Figure 3D**), and the maximum shell, also referred to as the maximum core (Alvarez-Hamelin, Dall’Asta et al. 2006), contained mostly associative ROIs involved in higher order cognitive function (**Figure 3E**). This bias towards associative ROIs subserving cognitive function within the maximum core occurred in ∼89% subjects (**Figure 3B**).

**Figure 2.**
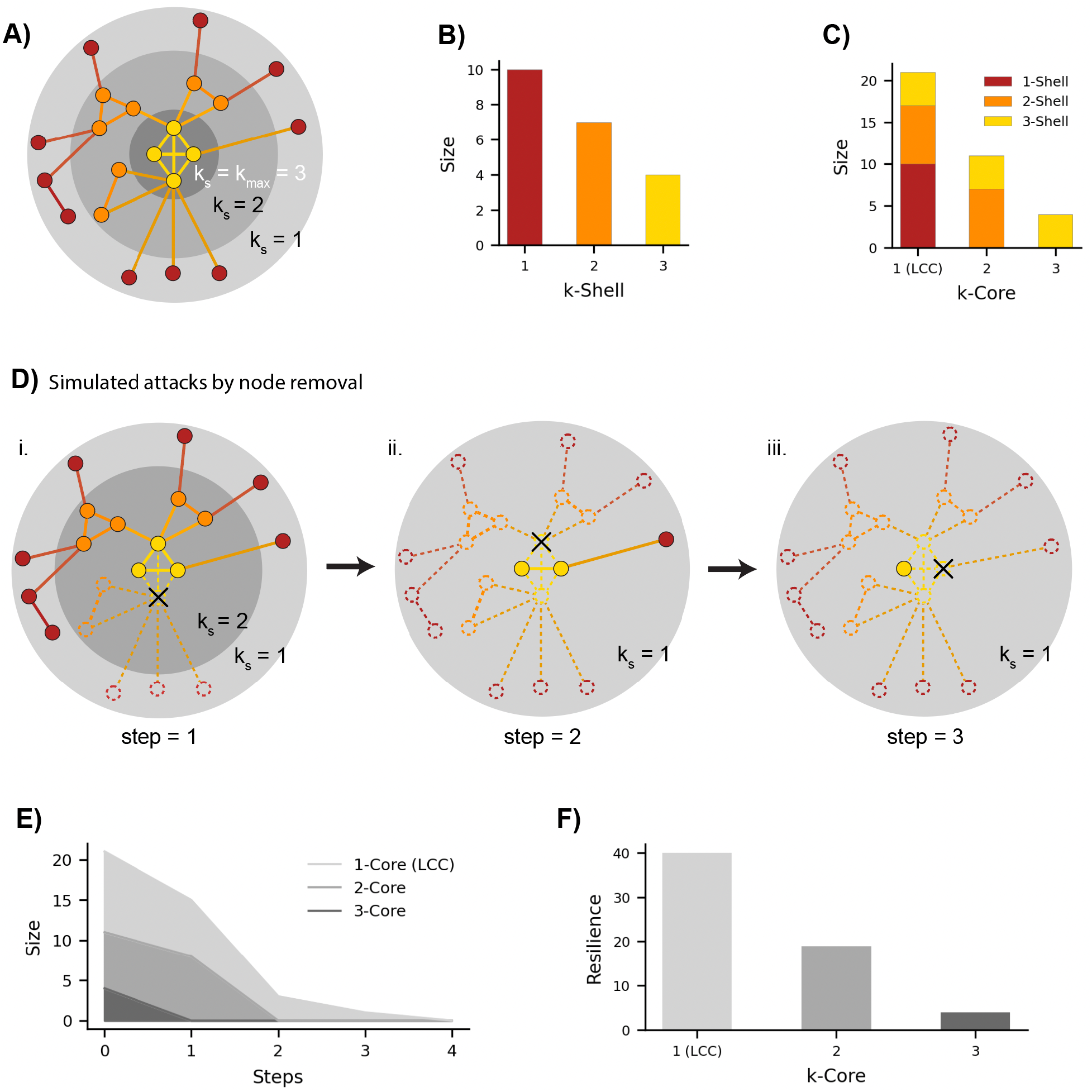
*k-*shell decomposition, and evaluation of network resilience. *k-*shell decomposition algorithmically decomposes networks into hierarchically organized shells of increasing connectivity. For a node to be placed in the k^th^ shell it must have degree k after removing all nodes with degree less than *k*. This process is iteratively repeated until only nodes with degree greater than or equal to k remain within the network. A node within the k^th^ shell is also within the k^th^ core. However, the k^th^ core also includes nodes in higher shells with degree greater than k. The maximum shell of a network is called the max core of the network and is denoted by k_max_, in the example above, k_max_ = 3. **A)** *k*-shell decomposition of a simple network before any attacks have been performed. The shell each node belongs to is denoted by their color, while the core is denoted by which concentric gray circle the node is contained within. **B)** Size of each *k*-shell in the example network. **C)** Size of each *k*-core, with colors denoting the shell memberships for each node. **D)** Simulating targeted node removal on the network for three time-steps (i, ii, iii). Nodes retain their original shell color during the simulation, however the *k*-shell (and k-core) they belong to decreases as nodes are removed. This is visually represented by the removal of the gray circles for 3-core in step 1 (i), and the 2-core in step 2 (ii). **E)** Size of each k-core as the simulation progresses. **F)** Quantification of resilience as the sum of core sizes at each time-step in the simulation. Resilience calculated for nodes and networks via shell assignment is similar, but instead we sum the *k* values that nodes are assigned at each time-step to get nodal resilience, to get *k*-shell resilience for networks, we averaged the nodal resilience for all nodes within a network.

**Figure 3.**
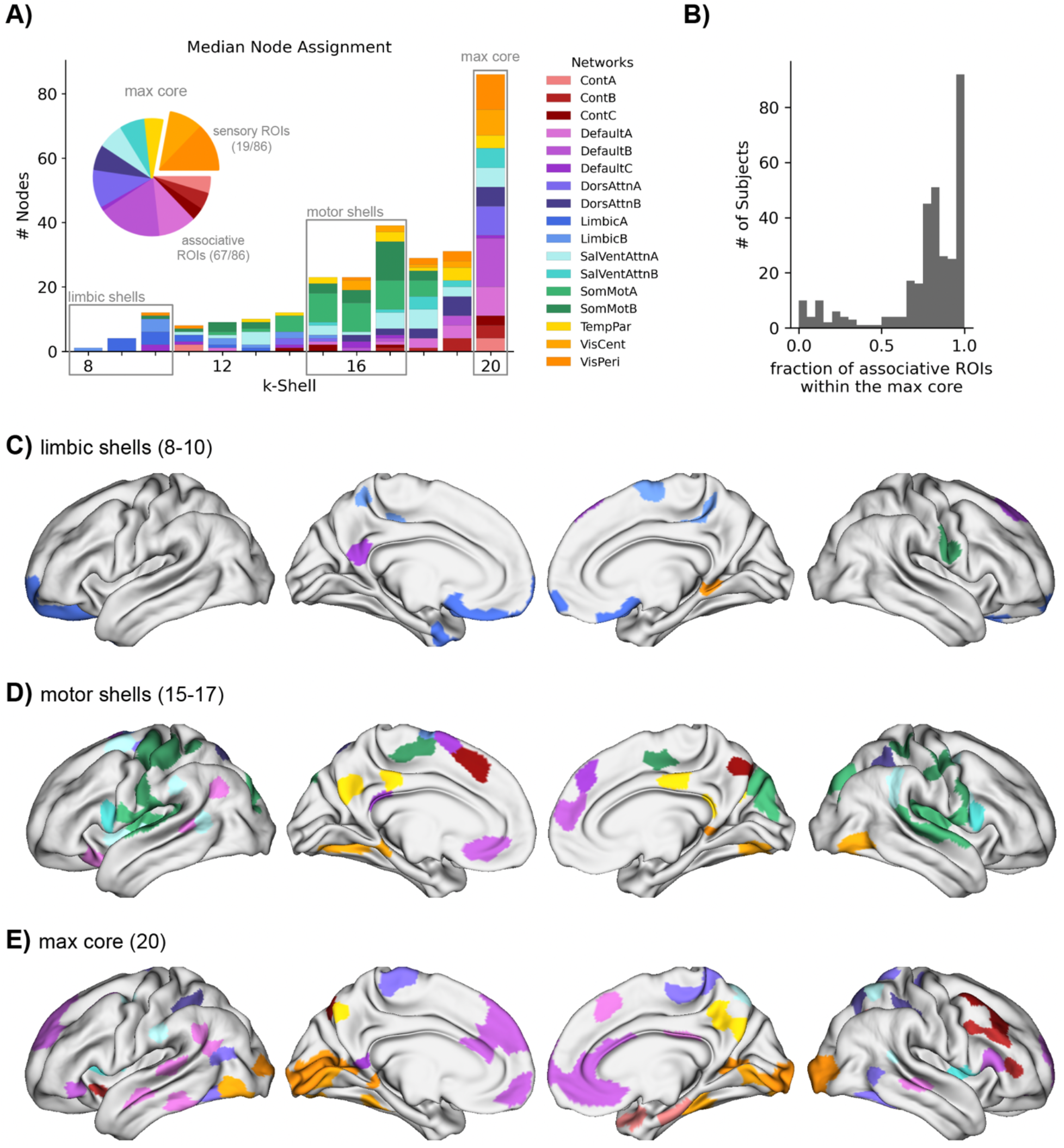
*k*-shell decomposition of brain networks. **A)** Median *k*-shell assignment for each region across subjects, with colors denoting which network nodes belong to. The cognitive core observed within the median *k*-shell assignment was mostly filled with associative ROIs (part of the FPCN, DMN, DAN, SVA, TP networks). **B)** Subjects tended to have more associative regions than sensory regions within their respective max core. **C)** Surface plots of lower *k*-shells primarily filled with nodes in the Limbic network **D)** Surface plots of mid-ranged Shells primarily filled with nodes in the Motor networks. **E)** Surface plot of the Cognitive Core observed in the median node assignment. (FPCN = Frontoparietal Control Network, DMN = Default Mode Network, DAN = Dorsal Attention Network, SVA = Salient Ventral Attention Network, TP = Temporal Parietal)

### 2.2 Episodic memory is related to the presence of robustly connected core nodes

We next investigated if the presence of robustly connected core nodes related to high cognitive performance during aging. We used episodic memory performance derived from the Rey Auditory and Verbal Learning Task (Rey 1941) as our measure of cognitive performance. This cognitive measure showed the expected negative correlation (r=-.248, p=7.6e-6) between age and task performance within our subjects (**Figure 4A**). In all following analyses, we included age as a covariate to identify effects that relate to high cognitive performance over and above subjects’ age. We next studied if episodic memory was related to the number of nodes within each *k*-core. As described above, a *k*-core contains all shells with value *k* or higher. As the value of *k* increases, nodes must be more robustly connected within the network in order to remain a part of it. The average number of nodes in each *k*-core decreased non-linearly as a function of *k* (r^2^=.941, p=2.418e-17) (**Figure 4B**). For lower values of *k*, we see relatively small differences between subjects. However, as the value of *k* increased, the variance in *k*-core sizes among subjects increased until the 21-core, at which point many subjects lacked these highly connected cores. The sizes of these cores were negatively correlated with participant age for cores 3-20, 25, & 26 (all p-values < 0.05) (**Figure S1B**), which illustrates the sensitivity of *k*-shell decomposition to age-related changes in functional connectivity. However, we found that the number of nodes within the *k*-cores median-high, were positively related to episodic memory performance (all p-values < 0.05) (**Figure 4C**).

**Figure 4.**
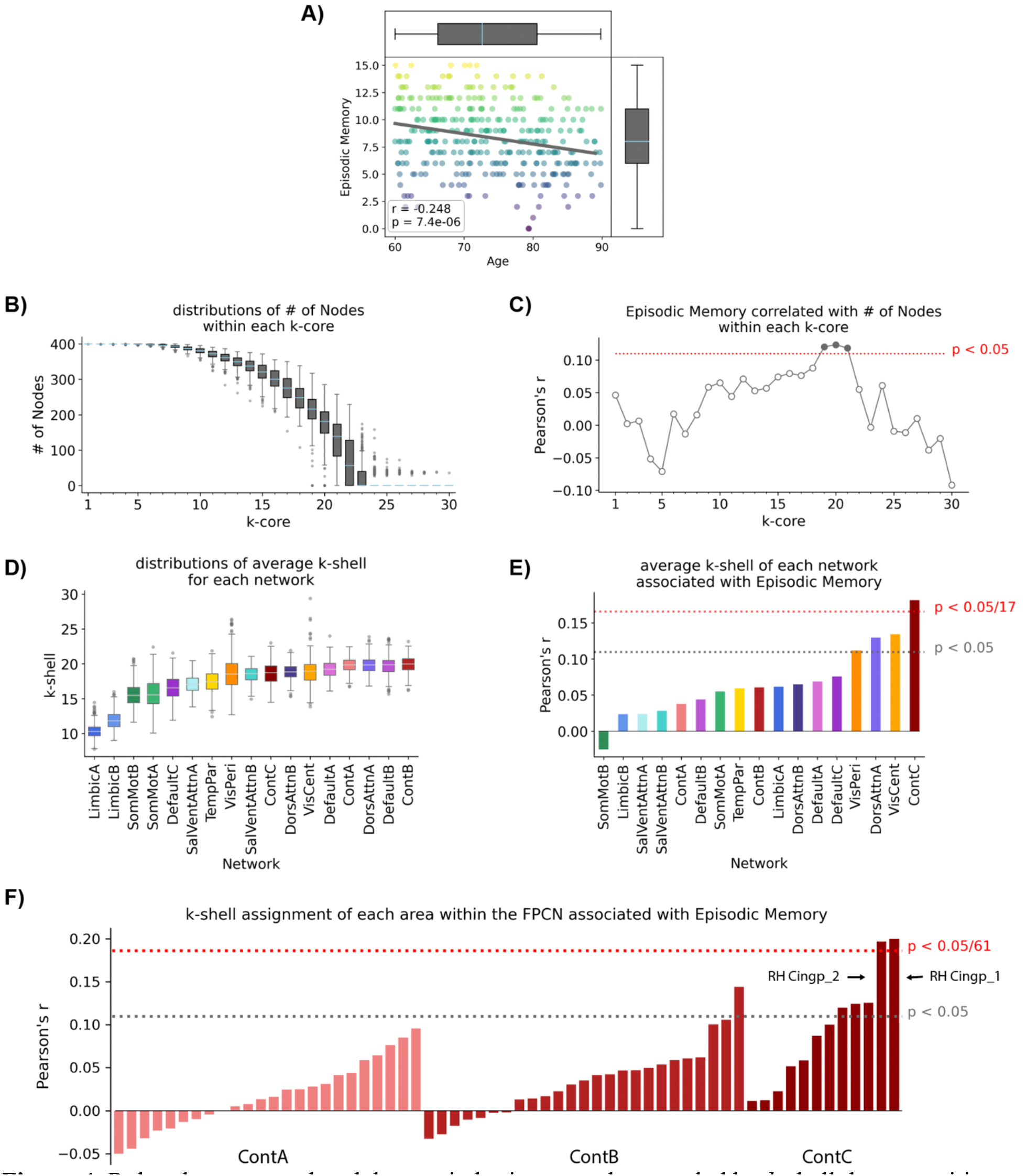
Robustly connected nodal cores in brain networks revealed by *k*-shell decomposition are associated with Episodic Memory. **A)** Episodic memory was negatively associated with age of participants, which was subsequently included as a covariate in all analyses involving episodic memory. **B)** Distributions of the size of each k-core among all subjects. **C)** Episodic Memory was significantly associated with the number of nodes within deep *k*-cores 19-21. **D)** Distributions for the average *k*-shell assignment per network after *k*-shell decomposition. **E)** Episodic Memory performance was positively associated with the average *k*-shell assignment of the Frontoparietal Control subnetwork – ContC (corrected p-value < 0.05). **F)** The *k*-shell assignment of two right hemisphere posterior cingulate regions within the Frontoparietal Control Network was related to Episodic Memory (corrected p-value < 0.05).

### 2.3 Episodic Memory is associated to robustly connected nodes within the Frontoparietal Control Network

Next, we investigated if the relationships between robustly connected nodes and Episodic Memory varied between networks. To do so, we calculated the average *k-*shell assignment for all nodes within each functional network and associated these measures with episodic memory. We found that the distributions of these averages followed similar patterns to those observed in the median shell assignment (**Figure 4D**). Specifically, the average *k-*shell assignment was the lowest for limbic networks, and highest for subnetworks within the Frontoparietal Control Network (FPCN), Default Mode Network (DMN), and the Dorsal Attention Network (DAN). As in the previous analysis with core size, the average *k*-shell assignments of 7/17 networks were negatively associated with age after using the Bonferonni method to correct for multiple comparisons (for 14/17 networks p-values < 0.05) (**Figure S1C**). When age was included as a covariate, we found that average *k*-shell assignment of the FPCN subnetwork – ContC and Episodic Memory were significantly correlated (r=0.181, p=0.0013) (**Figure 4E**).

On an individual ROI-level, the *k*-shell assignment of most ROIs within the FPCN were moderately related to episodic memory at best (**Figure 4F**). However, we found two right hemisphere posterior cingulate regions within ContC that were highly related to episodic memory (MNI coordinates: [(7, -44, 20), (6, -26, 28)] r=[0.200, 0.198], p-values =[0.0003, 0.0004], respectively) (**Figure 4F**). The median shell assignment of these two regions were 15, and 17, for the regions at MNI coordinates (7, -44, 20), and (6, -26, 28), respectively (**Figure S2C**), which are both larger than the size of the ContC network. This suggested that these areas might serve an integrative role among functional networks that promotes performance in the episodic memory task. To investigate this possibility, we identified regions within their *k*-shells outside of the ContC network. We found that an edge connecting one of the posterior cingulate regions (MNI coordinates: (7, -44, 20)) and a temporal parietal region (MNI coordinates: 59, -46, 7), was highly associated with episodic memory performance (ANCOVA, DF=15, F=5.92, p=8.71e-11, corrected p=7.00e-6).

### 2.4 The resilience of robustly connected core nodes is related to episodic memory

Following our analyses of the relationships between robustly connected core nodes and episodic memory, we sought to relate the resilience of these nodal cores to episodic memory. First, we measured resilience of cores within brain networks to targeted attacks. To do so, we rank nodes by degree, and then remove each node in descending order until no nodes remain (**Figure 2D**). After each node removal, we calculate the size of each *k-*core to track whether each particular core collapses or resists node removal (**Figure 2E**). Finally, we sum the size of each respective *k-*core across all time-steps to provide a single measure of resilience for each *k*-core (**Figure 2F**). This approach is similar to the one proposed previously (Schmidt, Pfister et al. 2019). However, our method enabled us to study the resilience of each core within a network, and we continued node removal until all cores have collapsed. As expected, *k*-core resilience measures varied as a function of *k* (r^2^=.996, p=6.35e-33), (**Figure 5A**). We note that these core resilience measures showed negative associations with age for cores 1-26 (all p-values < 0.05) (**Figure S3A**), as a result, we included age as a covariate when associating *k*-core resilience with episodic memory. We found positive relationships between resilience in cores 15-20 and episodic memory (all p-values < 0.05) (**Figure 5B**). Some of these relationships resembled our previous analysis that found correlations between the size of *k*-cores 19-21 and episodic memory (**Figure 4C**). These results suggests that robust core architectures may enable a form of network resilience, invariant to age, that supports episodic memory.

**Figure 5.**
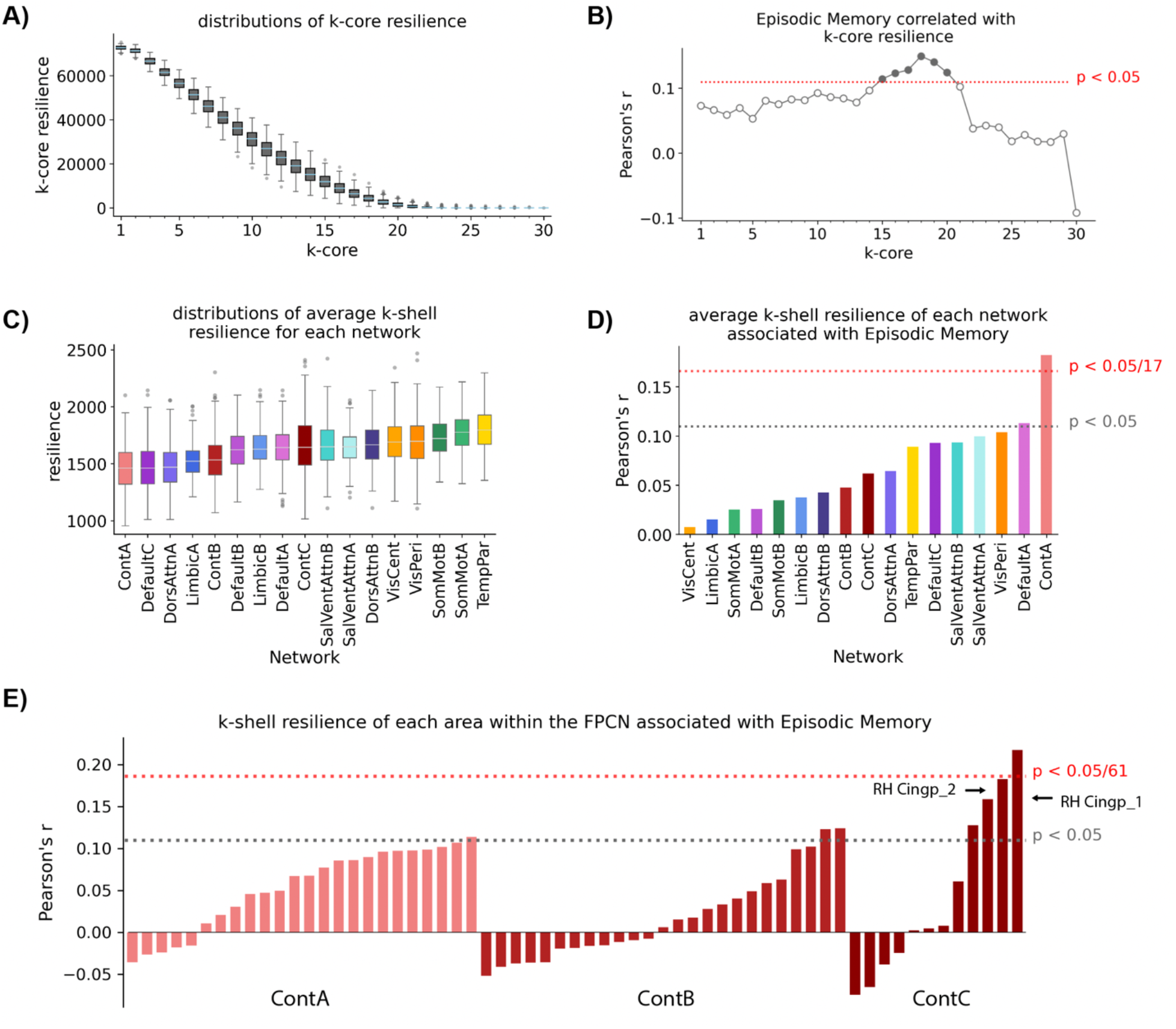
The resilience of robustly connected nodal cores is positively associated with Episodic Memory. **A)** Distributions of *k*-core resilience among all subjects. **B)** Resilience was positively associated with Episodic Memory for cores 15-20. **C)** Distributions for the average *k*-shell resilience per network. **D)** The *k*-shell resilience of the ContA network was positively associated with Episodic Memory. **E)** The *k*-shell resilience in a right hemisphere, posterior cingulate region within ContC showed a strong positive relationship with Episodic Memory (corrected p < 0.05). In all correlations performed above, age is included as a covariate.

### 2.5 Episodic Memory is associated to the resilience of robustly connected nodes within the Frontoparietal Control Network

Next, we related the *k*-shell resilience of each functional network to episodic memory. As done in the previous analysis (**Figure 4D**), we calculate the average *k*-shell assignment of each functional network, and then perform targeted attacks, recalculating the average *k*-shell assignment for each functional network after each attack. We summed these values across all time steps to give a single measure of *k*-shell resilience for each functional network. These measures give an indication of each network’s ability to maintain robust connectivity despite perturbation. We found that *k*-shell resilience varied across the different networks (**Figure 5C**). As before, we associated these measures with age and found negative relationships between *k*-shell resilience of functional networks and episodic memory (all p-values < 0.05, 14/17 corrected p-values < 0.05) (**Figure S3B**). When we included age as a covariate, we found a significant positive relationship with a FPCN – ContA resilience and episodic memory (r=0.182, corrected p=0.0187) (**Figure 5D**).

We next moved from the network level down to investigating individual ROIs within the FPCN. For a single ROI, we measured the *k*-shell resilience by summing the *k*-shell assignment of the ROI across all time-steps of the attack simulations. Similarly to our measure of *k*-shell resilience for networks, these resilience measures provide an indication of an individual ROI’s ability to maintain robust connectivity to the larger network despite perturbation. Within the FPCN we associated the *k*-shell resilience of each ROI to episodic memory (**Figure 5E**). We found that the *k*-shell resilience of one of the same right hemisphere posterior cingulate regions within ContC found in the previous analysis (MNI=(7, -44, 20)) (**Figure 4F**) was significantly related to episodic memory (r=0.217, corrected p=0.0056) (**Figure 5E**). Despite observing a significant positive relationship between average *k*-shell resilience of the ContA subnetwork and episodic memory, the resilience of all individual ROI within ContA were not significantly correlated with episodic memory (all corrected p-values > 0.05).

### 2.6 The presence and resilience of robustly connected nodes is not associated with Processing Speed

We tested if the relationships we discovered thus far were specific to episodic memory or rather extended to other cognitive functions. To do so, we related topological features and resilience measures with processing speed rather than episodic memory (**Figure S4**). We did not observe similar relationships between processing speed and *k*-core size (**Figure S4B**), k-core resilience (**Figure S4C**), average *k*-shell of each network (**Figure S4D**), or average *k*-shell resilience of each network (**Figure S4E**). As before, age was included as a covariate in all analyses.

### 2.7 Segregation of FPCN – ContC mediates relationships between network resilience and Episodic Memory

The results reported above suggest that robust connectivity and resilience of nodal cores involving the FPCN are both related to episodic memory. Network segregation (**Figure 6A**), a widely-used quantification of segregation in the brain, measuring differences in connectivity within, relative to between networks (Chan, Park et al. 2014, Wig 2017, Chan, Han et al. 2021), has been widely linked to cognitive performance in aging (Chan, Park et al. 2014, Wig 2017, Chan, Han et al. 2021) and to cognitive resilience in AD (Ewers, Luan et al. 2021). In particular, segregation of the FPCN has been associated with greater episodic memory in aging (Geerligs, Renken et al. 2015). Our results, combined with these previous findings, motivated us to investigate if the associations between resilience of nodal cores and episodic memory were mediated by functional network segregation. We quantified the correlation between segregation in each functional network and k-core resilience and found widespread relationships with many of the individual networks (**Figure 6B**). We also evaluated the extent to which the association between segregation of each functional network and episodic memory persisted, over and above age (**Table S1**). In our sample, only the correlation between segregation of the FPCN - ContC was significantly correlated with episodic memory, irrespective of age (**Figure 6C**). Lastly, we performed a parallel mediation analysis to test if segregation of any of the functional networks in our parcellation mediated relationships between resilience of nodal cores and episodic memory (**Figure 6D**). We found that FPNC - ContC segregation mediated relationships between *k*-core resilience and episodic memory for cores 1-15 (corrected p-values < 0.05) (**Figure 6E**).

**Figure 6.**
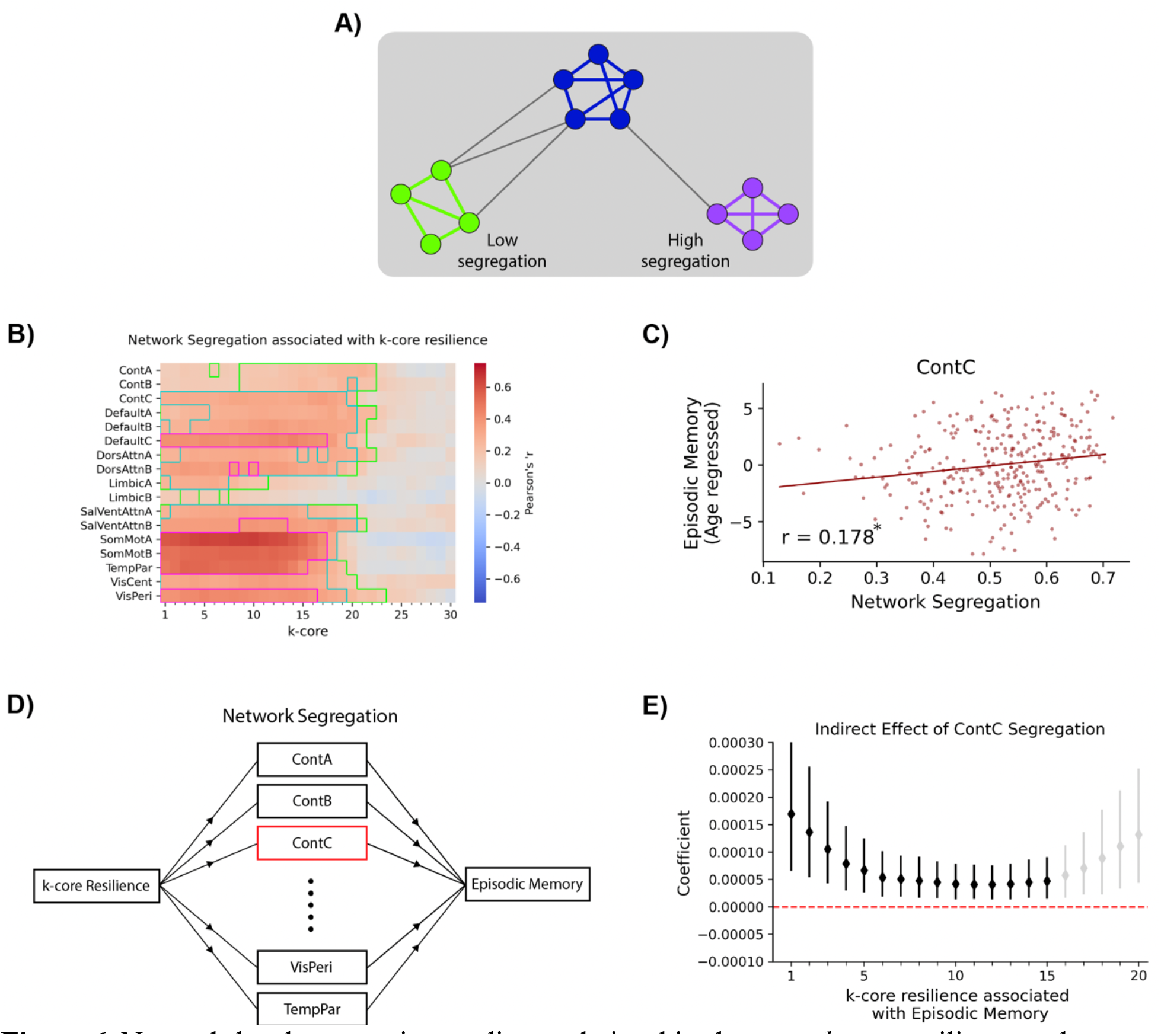
Network-level segregation mediates relationships between *k*-core resilience and Episodic memory in aging. **A)** A toy network illustrating the differences between networks with low (green) and high (purple) segregation. **B)** Segregation of each subnetwork associated with *k*-core resilience. Areas on the heatmap with corrected p-values < 0.05, 1e-5, and 1e-10, are bounded by the green, cyan, and magenta borders, respectively. **C)** Segregation of ContC network was significantly associated with Episodic Memory after controlling for the effects of age and education, in addition to performing correction for multiple comparisons (corrected p-value < 0.05). **D)** We performed a serial mediation to examine if network segregation of any of our 17 networks mediate relationship(s) between *k*-core resilience and Episodic Memory. **E)** The indirect effect of ContC segregation was significant for cores 1-20, however, after correcting for multiple comparisons (corrected p-value < 0.05), this relationship only remained for *k*-cores 1-15 (denoted in black). Subject age is included as a covariate in the correlations and mediation analyses performed above.

### 2.8 Relationships between robustly connected nodes and redundant functional short paths

We next examined whether robustly connected nodes tended to exhibit a relatively increased number of shortest paths, a topological property that may increase nodal redundancy (De Vico Fallani, Rodrigues et al. 2011), in-itself a protective mechanisms in cognitive aging (Langella, Sadiq et al. 2021). We found that the measures of robust connectivity generated from *k*-shell decomposition are generally related to the number of 1, 2, and 3-step paths between nodes within the same shell (**Figure S5**). Specifically, as *k*-shell assignment of a node increases in the value of *k*, there was generally an increase in the number short paths between nodes within the same shell.

### 2.9 Sensitivity to parameter selection and additional controls

The process for generating binarized networks from measures of RSFC in this study required two parameter choices, the number of OMSTs, and the frequency threshold that we used to determine what edges we kept (**methods**). We repeated the analyses in the main results of our study across several different parameter choices. These choices impacted the density of our networks (**Figure S6**). However, the previously identified patterns were generally consistent between parameter selections. In particular, robust connectivity and resilience of robustly connected nodes were both associated with episodic memory (**Figure S7**). The similarities of the significant results between these two analyses prompted us to perform an additional experiment that included the core size as a covariate when correlating resilience with episodic memory, which did not yield any significantly different results (**Figure S8**). Finally, FPCN segregation consistently mediated relationships between core resilience and episodic memory (**Figure S9**). We refer readers to our **supplemental results** for more information.

## 3. Discussion

In the current study we examined the relationships between the robust core architecture of the human brain, network resilience, and episodic memory in healthy aging. We found that episodic memory performance was positively related with robust core connectivity structure, particularly involving the FPCN. Many of these same nodal cores exhibited resilience to targeted attack simulations that was also related with episodic memory. We found that segregation of a subnetwork within the FPCN (ContC) mediated relationships between resilience of nodal cores and episodic memory. We additionally demonstrated that the observed findings are specific to memory performance, and do not extend to other cognitive functions, such as processing speed.

### 3.1 Robust connectivity and resilience of nodal cores both promote episodic memory in aging

Nodes within the higher *k*-cores and *k*-shells of networks have been found to play an important role in their stability (Kitsak, Gallos et al. 2010, Berry and Widder 2014, Zhang, Zhang et al. 2017, Morone, Del Ferraro et al. 2019, Burleson-Lesser, Morone et al. 2020). While we do not make a direct comparison between sizes of *k*-cores and *k*-shells to network resilience, we did find that having more nodes in higher *k*-cores is associated with greater cognitive function. We also found relationships between network resilience and episodic memory in similar subsets of cores to those that possessed the aforementioned relationships. In particular, we found that nodes within the FPCN displayed more robust connectivity and greater resilience in individuals who performed well on the episodic memory task. These findings indicate that, as in other types of complex systems (Kitsak, Gallos et al. 2010, Berry and Widder 2014, Zhang, Zhang et al. 2017, Morone, Del Ferraro et al. 2019, Burleson-Lesser, Morone et al. 2020), core network structure is an important component in promoting network resilience, promoting intact cognitive function in healthy aging.

### 3.2 The FPCN in health and disease

Our results demonstrate a specialized role for the FCPN in robust network connectivity and resilience. These findings are in strong agreement with earlier reports. The FPCN has been shown to play an important role in cognitive function in healthy aging and in disease. In the context of healthy aging, greater functional segregation between the FPCN and DMN has been associated with greater verbal episodic memory (Geerligs, Renken et al. 2015). Additionally, decreased FPCN segregation has been shown to relate to poorer executive function (Sims, Faulkner et al. 2021) and global cognitive performance (Chong, Ng et al. 2019). Longitudinally, declines in FPCN segregation have been associated with reduction in processing speed (Malagurski, Liem et al. 2020). Other studies have implicated FCPN connectivity in age-associated disease states. For example, an increased coupling of the FPCN and DMN has been shown in Alzheimer’s disease (Contreras, Avena-Koenigsberger et al. 2019).

Our results point to an association between FPCN segregation and verbal episodic memory in healthy aging, consistent with previous findings (Geerligs, Renken et al. 2015). However, using attack simulations, we find that FPCN segregation mediates relationships between network resilience and episodic memory. These results support the notion that segregation in brain networks might provide a form of cognitive ‘reserve’ that slows cognitive decline (Wig 2017, Chan, Na et al. 2018), as has recently been demonstrated in Alzheimer’s disease (Ewers, Luan et al. 2021). In addition, we found that resilience of the FPCN was positively related to episodic memory, which aligns well with the recently discovered association between FPCN resilience in Parkinson’s disease and protection against cognitive decline (Cascone, Langella et al. 2021).

### 3.3 Robust connectivity and redundancy in functional networks

We found that robust connectivity within nodal cores and the FPCN were both positively related to episodic memory. One possible reason for this association is that more robust connectivity could provide a greater number of redundant functional pathways leading to the activation and/or maintenance of activity in brain regions responsible for task performance. Previous studies have indicated a role for redundancy within the hippocampus in mediating relationships between hippocampal volume and memory (Langella, Mucha et al. 2021), demonstrated that redundancy within the hippocampus is lower in individuals with mild cognitive impairment (MCI) (Langella, Sadiq et al. 2021), and found that redundancy was related to greater executive function in aging individuals (Sadiq, Langella et al. 2021). The positive relationship between robust connectivity, as determined by *k*-shell decomposition, and number of short paths between nodes along with robust connectivity correlating with episodic suggests that redundancy in core network structure and within the FPCN may play a role in facilitating episodic memory in healthy aging.

### 3.4 Comparisons to other methods for evaluating network resilience in brain networks

The most commonly employed method for studying network resilience in brain networks is to perform node removals and evaluate the ability of the network to remain connected. This is done by calculating the fraction of nodes within the largest connected component (LCC), before and after each simulated attack have been performed and then calculating the area under the curve (AUC) for the fractional LCC values at each timestep (Albert, Jeong et al. 2000, Piraveenan, Thedchanamoorthy et al. 2013). Attack simulations utilizing LCC resilience have been used to study changes in network resilience associated with neurodegenerative (Mancini, De Reus et al. 2016, Cascone, Langella et al. 2021) and neuropsychiatric (Palaniyappan, Hodgson et al. 2019) diseases. In our analyses, the 1-core, all connected nodes with at minimum degree 1, is equivalent to the LCC. Therefore, values we obtained for the resilience of the 1-core are mathematically equivalent to those that would be obtained by measuring the AUC for fractional LCC values during simulated attacks if we normalize by the starting size of the 1-core. Since this normalization results in a linear transformation of the data, the association and mediation analyses we performed will be unaltered. Additionally, since the starting size of a *k*-core is an import feature that relates to the resilience of the *k*-core (Burleson-Lesser, Morone et al. 2020), we maximize the opportunity to observe those relationships in our data by avoiding normalization for all *k*-cores. However, we repeated the partial correlations between core size and episodic memory with starting core size for each *k*-core as a covariate when studying *k*-core resilience, and found that the results were largely consistent between the two methods.

We found that resilience of the LCC for functional brain networks is mostly unrelated to episodic memory for healthy aging subjects across a wide variety of parameter settings (**Figure S6D**).

The methods we used for studying core resilience and changes in shell assignment at nodal and functional network levels provided more sensitivity to differences in robust connectivity profiles, which enabled us to find relationships between measures of resilience and episodic memory. While studying core resilience has been done before in financial markets (Burleson-Lesser, Morone et al. 2020) and ecological networks (Morone, Del Ferraro et al. 2019), this method has not been applied towards understanding network resilience in the brain.

### 3.5 Between-subject variability during network construction

Construction of functional brain networks using OMST-based methods tends to increase the amount of between-subject variability when applied to functional brain networks (Jiang, Betzel et al. 2021). However, our analyses comparing different parameter selections showed that for the network features we studied, the strongest relationships with episodic memory occurred when we reduced the variability in initial network construction (via thresholding edges by their frequencies) (**Figure S7**). These relationships tended to peak when we kept edges only if they occurred at the 75^th^ quantile of edge frequencies among all subjects. One possible implication of this result is that only ∼25% of topological variability associated with OMST-based network construction is needed to find topological features relating to differences in episodic memory performance during aging. However, a related explanation could be that the increased between-subject variability in network structure decreases the signal-to-noise ratio when associating topological features to behavioral traits such as episodic memory. Future studies will be needed to untangle the behavioral relevance of the additional variability associated with OMST-based network construction.

### 3.6 Limitations

For network construction, we used an OMST-based method. We tested a large set of parameters and observed consistent results across parameter settings. However this process requires more choices from the user than a data driven OMST-based network construction process like the one proposed previously (Dimitriadis, Antonakakis et al. 2017). As a result, it’s possible that the parameters we sampled led to us to missing other potential relationships within our dataset. However, the consistency we observed across parameter settings indicated the robustness of our results, and aligned well with the existing literature.

## 4. Conclusion

Using graph analysis methods we decomposed brain networks into hierarchically layered shells, finding that robust connectivity in nodal cores, particularly in the FPCN, support superior cognitive performance in healthy aging. Additionally, we use attack simulations to study the resilience of these nodal cores and find that resilience, particularly in the FCPN, is strongly related to episodic memory in healthy aging. Finally, we build on existing literature studying the role of brain segregation in cognitive function by demonstrating that segregation within the FPCN mediates relationships between network resilience and episodic memory. Altogether, the results highlight the importance of robust and resilient functional networks in healthy cognitive aging.

## 5. Methods

### 5.1 Dataset and participants

Data were obtained from the Human Connectome Project – Aging database (HCP-Aging) (Harms, Somerville et al. 2018), part of the 2.0 Release of the data. Subjects ranged from the ages of 60 – 90 (Female/males= 171/149) and demonstrated normal cognitive function as assessed by the Montreal Cognitive Assessment. For subjects 60-69 years of age, normal cognitive function was a score greater than or equal to 26, whereas for subjects older than 70, their scores needed to be within one standard deviation of their age and education adjusted norm (Malek-Ahmadi, Powell et al. 2015). In total, 320 subjects with available resting-state functional magnetic resonance imaging (fMRI), structural MRI, and cognitive measures were used. All subjects provided written informed consent and all procedures were approved by the local Institutional Review Boards

### 5.2 Image processing

A 3 Tesla Siemens Prisma scanner was used to collect functional and structural images (Harms, Somerville et al. 2018). A multi-echo magnetization prepared rapid gradient echo (MPRAGE) sequence (voxel size: 0.8×0.8×0.8mm, T =1.8/3.6/5.4/7.2ms, TR=2500ms, flip angle=8 degrees) was used to collect structural images. Functional imaging was collected using a 2D multiband gradient-recalled echo echo-planar imaging sequence (voxel size: 2×2×2mm, TE=37ms, TR=800ms, flip angle=52 degrees). Two functional scans were taken for each session with opposite phase encoding directions (poster-anterior, anterior-posterior). Subjects were instructed to keep their eyes open on a fixation cross during functional scans. Structural and functional images underwent minimal preprocessing according to the HCP-pipeline, which included spatial artifact/distortion removal, cross-modal registration, and alignment to standard space (Glasser, Sotiropoulos et al. 2013). Additional image processing was done in the Matlab package CONN (Whitfield-Gabrieli and Nieto-Castanon 2012), which included outlier identification (movement of more than 0.9mm, or global blood-oxygenation dependent signal changes greater than 5 standard deviation of the mean), nuisance regression (white matter, cerebrospinal fluid, and 12 motion parameters), and band-pass filtering at 0.033 – 0.083 Hz (Buzsáki and Draguhn 2004). Functional timeseries were obtained for the Schaefer Local-Global 400 regions of interest (ROI) – 17 network parcellation (Schaefer, Kong et al. 2018), which is a cortical parcellation that encompasses cognitive, sensorimotor, and visual regions. 400 × 400 Fisher Z transformed correlation matrices were calculated from these timeseries for each participant.

### 5.3 Network construction

We used a two-step process for constructing networks from each subjects resting state functional connectivity matrix (**Figure 1B**). The first step involves selecting *N* orthogonal minimum spanning trees (OMSTs), similar to the strategy proposed before (Dimitriadis, Antonakakis et al. 2017). For each minimum spanning tree (MST) (Meier, Tewarie et al. 2015), this is done by constructing a tree that connects every node within the graph with fewest possible edges and the largest possible sum of edge weights. Each tree constructed will be used for the final network, but all subsequent trees must be built without using any edges from previous trees. After constructing *N* trees, they are fused together for the final network. In the second step, we filter edges to reduce the variability of the networks generated. The filter was determined by calculating the frequency of each edge pair among all subjects, and then keeping edges only if they occurred at the *X*^th^ quantile of edge frequencies. This is formally defined bellow.

1. Let *A*_*r*×*r*×*n*_ be the group binary adjacency matrix constructed from, where *r* is the size of the parcellation, and *n* is the number of subjects.
2. Then, for each *k, A*_:,:,*k*_ refers to the binary adjacency matrix constructed from *N* OMSTs for the *k*^*th*^ subject.
3. Now, *F*_*r*×*r*_ is the frequency matrix where each *F*_*i,j*_ denotes the frequency of occurrence of an edge between nodes *i* and *j* in the group matrix *A*_*r*×*r*×*n*_. For binary matrices, this can be calculated by taking the mean across subjects.
4. Calculate the frequency threshold *θ* by taking the *X*^th^ quantile of the frequency matrix *F*_*r*×*r*_:

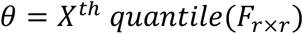
5. Finally, step through all subjects *k*, and node pairs (*i, j*), in each subject’s binary adjacency matrix *A*_:,:,*k*_, and only keep an edge if *F*_*i,j*_ > *θ* and *A*_*i,j,k*_:

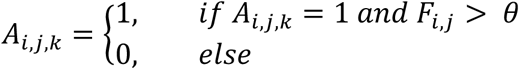

We sampled several parameter combinations for *N*, the number of trees, and *X*, the quantile of the frequency at which we thresholded. The analysis that follows is focused on a representative parameter selection of 20 trees with edges thresholded at the 50^th^ quantile of edge frequencies. This approximately corresponds to an average network density of ∼8%. We note that our method for network construction has the drawback of parameter selection as opposed data driven methods (Dimitriadis, Antonakakis et al. 2017). However, we found that the networks generated by the data-driven OMST algorithm produced networks with uniform degree distributions, which prevented us from studying how variations in robustly connected nodal cores relate to episodic memory. To ensure that our results were consistent across a wide range of possible parameter combinations, we replicated the main analyses for several other selections of trees and frequency-based edge thresholds (**Supplemental Information**).

### 5.4 Cognitive measures

We use two cognitive measures in our study, episodic memory, and processing speed. Episodic memory scores were derived from subject performance on the Rey Auditory and Verbal Learning Task (RAVLT) (Rey 1941), which has been shown to decline with age (Leal and Yassa 2015). Participants were tasked with learning and recalling list A, which consisted of 15 semantically independent words. Subjects did this over the course of 5 trials. On the 6^th^ trial, a distractor list was presented and subjects were tasked with recalling it once. Afterwards, subjects were tasked with immediately recalling list A (immediate recall), and recalling list A again 20-30 minutes layer (delayed recall). Subject performance on the delayed recall was used as the episodic memory score for our study. For processing speed, were subjects asked to judge whether images shown next to each other were identical as quickly as possible. They were given 85 seconds to respond to as many image pairings as possible.

### 5.5 k-shell decomposition and k-core size calculations

k-shell decomposition partitions a network into hierarchically organized shells of increasing connectivity (**Figure 2A**). To calculate a network’s *k*-shell decomposition, the procedure starts at the value 1, for which all nodes with degree less than or equal to 1 and their connections are removed from the network. After the initial removal, there might be new nodes with degree less than or equal to 1. This removal process is repeated until only nodes with degree greater than 1 remain within the graph. These removed nodes make up the 1-shell. This procedure is then repeated for each value of *k* until all nodes within the network have been assigned to a value for *k*, which refers to the shell they belong to. When we refer to the k^th^-core of a network, this is a reference to all nodes assigned a value for their respective *k*’s that is greater than or equal to *k*.

### 5.6 Attack simulations and resilience calculations

Attack simulations were performed using modified scripts from the python package tiger (Freitas, Yang et al. 2021). Briefly, each network was passed to tiger, where nodes were ranked by degree, the number of other nodes a node is connected to directly by an edge, and then removed in descending order. At each step in the attack simulation, each network’s *k-*shell decomposition (**Figure 2A & 2D**) was re-calculated using the python package NetworkX (Hagberg, Swart et al. 2008). The first resilience metric we calculated, *k*-core resilience, denotes the ability of nodes to stay within a particular core during the attack simulations. This is done by measuring the size of each core at each time-step (**Figure 2E**) and summing across time-steps to give a single value for resilience for each respective core (**Figure 2F**). For instance, in **Figure 2A** we see that the size of the 2-core is 11 at step 0, 8 at step 1, and 0 at step 3. This gives a resilience score of 11+8+0 = 19 for the 2-core. *k*-core resilience calculated in this way is a monotonically decreasing value for all cores. We also measure the resilience of *k*-shell assignment associated with each ROI during the attack simulations. This is done by calculating each ROI’s shell assignment at each time-step in the attack simulation and then summing across all time-steps. Finally, we measure the *k*-shell resilience of each network by averaging the ROI resilience measures for each ROI within a network.

### 5.7 Brain system segregation

Brain system segregation is a method for measuring how segregated the correlated activity within a functional network is in comparison to its correlated activity between itself and other networks (Chan, Park et al. 2014, Wig 2017, Chan, Han et al. 2021). The formal definition is as follows:

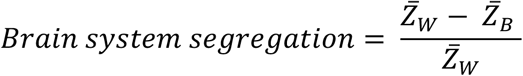

Where 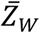 represents the mean within-systems correlation and 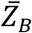 represents the mean between-systems correlation. We used brain system segregation to study relationships between network resilience and episodic memory (**Figure 1D**).

### 5.8 Statistical analysis

We performed correlations between network features studied here and age, and partial correlations between network features studied and episodic memory, while including age as a covariate. We also performed parallel mediation analyses between network resilience and episodic memory with the measures of segregation for each functional network as the potential mediators. The significance threshold set for these analyses was *α* = 0.05, and the Bonferonni method to correct for multiple comparisons was applied when appropriate. For test that considered each network separately this required dividing the significance threshold by 17. For tests considering the individual ROIs within the FPCN *α* was divided by the number of ROIS (61). We used a linear regression to plot the age-regressed relationship between episodic memory and segregation of FPCN-ContC. All statistical analyses were done using custom scripts in python. Mediations and partial correlations were performed with the python package Pingouin (Vallat 2018). Linear regressions were performed using SciPy (Virtanen, Gommers et al. 2020). Non-linear fits for the size and resilience of *k*-cores were calculated with Statsmodels (Seabold and Perktold 2010).

### 5.9 Plotting

Plotting and data visualization were done using custom scripts in python that made of use of the Matplotlib (Hunter 2007), Pandas (McKinney 2010), Seaborn (Waskom 2021), and Brainspace (de Wael, Benkarim et al. 2020) packages.

## Supporting information

Stanford_SI

## Author contribution

W.S., P.M., and E.D developed research plan; W.S. developed methods and analyzed data; W.S., P.M., and E.D interpreted results and wrote the paper.

