## Supplementary material for "A robust core architecture of functional brain networks supports topological resilience and cognitive performance in aging": Stanford_SI

### 1. Supplemental Results

*1.1 Sensitivity to parameter selection of network features associated with episodic memory*

As discussed in the **methods**, our analyses thus far focused on networks obtained from a single parameter selection which consisted of us constructing 20 OMSTs per subject, followed by thresholding edges that occurred at frequencies less than 50$^{th}$ quantile of edge frequencies among the entire group. Following our initial analyses, we analyzed how our main results would change given other parameter selections. We tested networks constructed with 10, 20, 30, and 40 OMSTs, with edges thresholded if they occurred at a frequency less than the 0$^{th}$, 25$^{th}$, 50$^{th}$, 75$^{th}$, and 95$^{th}$ quantile of edge frequencies among all subjects, for 20 combinations of parameters total. We illustrate how these parameter settings impact network densities in **Figure S6**. Associations between *k*-core size episodic memory across parameter settings were consistently higher in cores with more robust connectivity and parameter settings with reduced variability of edges kept (**Figure S7A**). The average shell assignment of FPCN-ContC was strongly associated with episodic memory across parameter settings (**Figure S7B**). The strength of the association between shell assignment of the right hemisphere posterior cingulate region (MNI coordinates: (7, -44, 20)) and episodic memory increased as we reduced variability of edges kept (**Figure S7C**).

*1.2 Sensitivity to parameter selection for resilience metrics associated with episodic memory*

The relationships between core resilience and episodic memory across parameter settings tended to reflect those observed with core size (**Figure S7D**). This observation led us to perform an additional control experiment where we calculated core resilience but including initial node count within each core as a covariate (**Figure S8A**). The z-scores for the differences before and after inclusion of this covariate were not significant for any of the parameters tested (**Figure S8B**). However, it is visibly clear that there are a reduced number of significant associations in **Figure S8A** when including core size as a covariate. The relationship between shell resilience of the ContA network and episodic memory was the most sensitivity to parameter selection, with only 6/20 parameter combinations retaining significance (**Figure S7E**). The association between episodic memory and shell resilience of the right hemisphere posterior cingulate region (MNI coordinates: (7, -44, 20)) was consistent across parameter settings, but tended to be stronger with more OMSTs and reduced edge variability (**Figure S7F**). Finally, we found that ContC segregation consistently mediated relationships between resilience of early cores and episodic memory across parameter settings (**Figure S9**). However, these indirect relationships tended to be the strongest when thresholding edges if they occurred at a frequency less than the 25$^{th}$ quantile of edge of edge frequencies among all subjects.

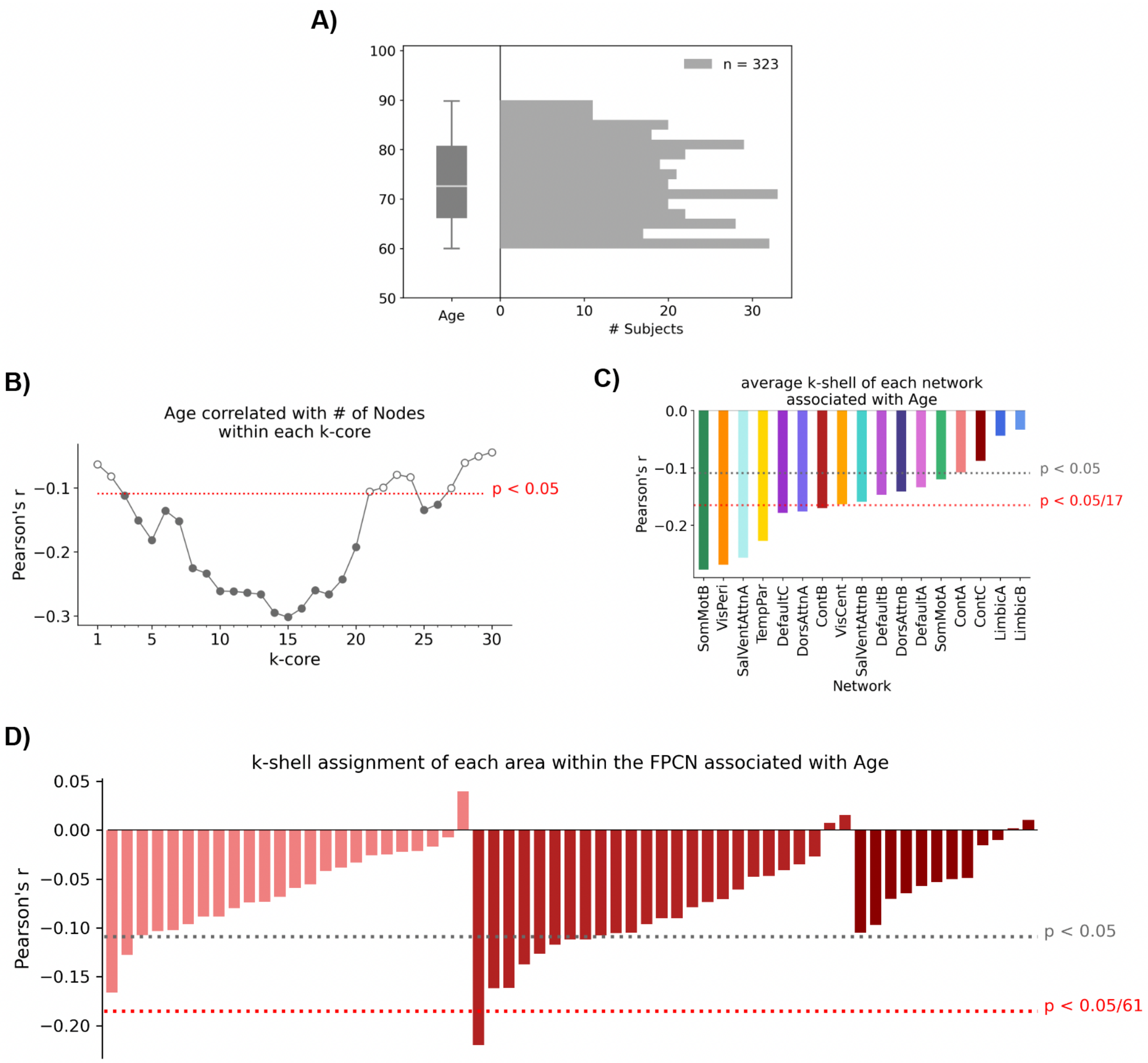


**Figure S1.** The presence of robustly connected nodal cores is negatively associated with age. **A)** Our subjects ranged in age from 60-90 years old. **B)** Age was negatively correlated with core size in cores 3-20, and 25-26. **C)** The average *k*-shell assignment of almost all networks were marginally associated with age in the negative direction, with several significant relationships remaining after corrected for multiple comparisons (corrected p-value < 0.05). **D)** The *k*-shell individual ROIs belonged to predominately showed negative correlations with age, with 20 remaining significant after correcting for multiple comparisons (corrected p-value < 0.0).

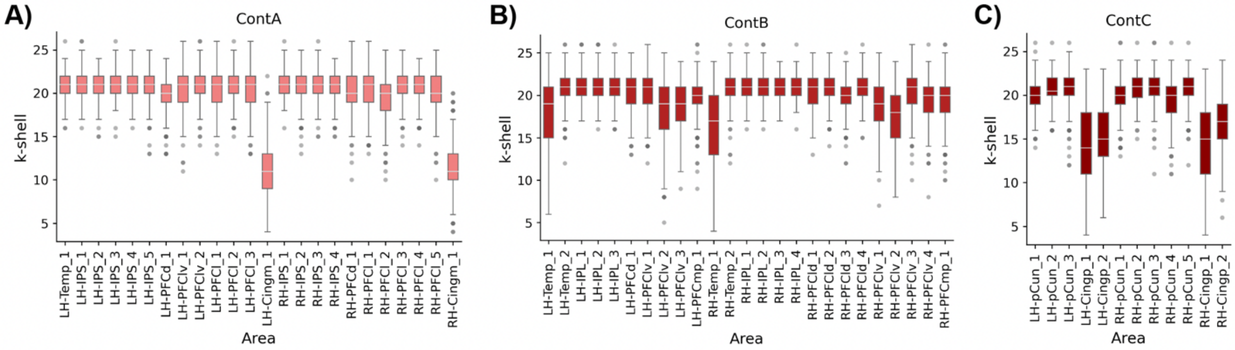


**Figure S2.** Distributions of *k*-shell assignments for ROIs within subnetworks of the Frontoparietal Control Network (FPCN). Subnetworks shown are **A)** ContA, **B)** ContB, and **C)** ContC.

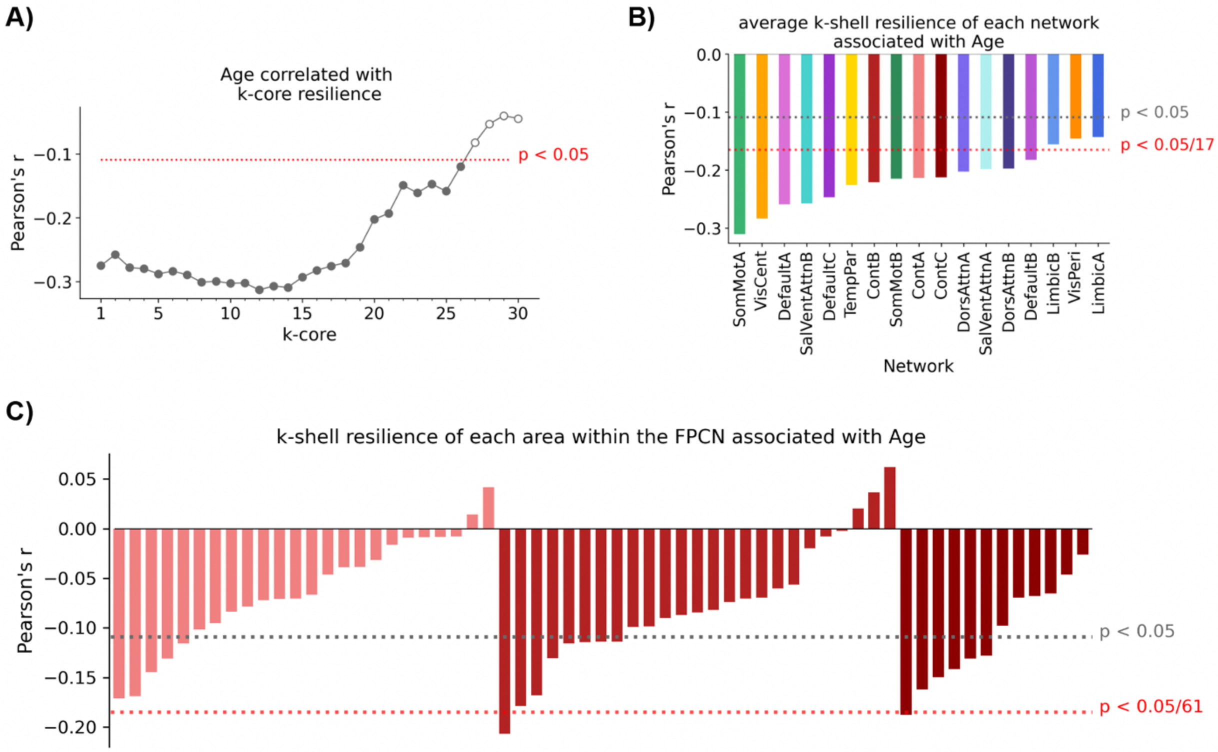


**Figure S3.** Age is associated with less resilient brain networks. **A)** The resilience against targeted attacks of almost every core examined was negatively associated with age. **B)** The average *k*-shell resilience of all but three networks were negatively associated with age. **C)** The *k*-shell resilience of individual ROIs within the FPCN were also generally negatively associated with age, with two ROIs, one within ContB, and one within ConC, showing significant negative relationships after correcting for multiple comparisons (corrected p-values < 0.05).

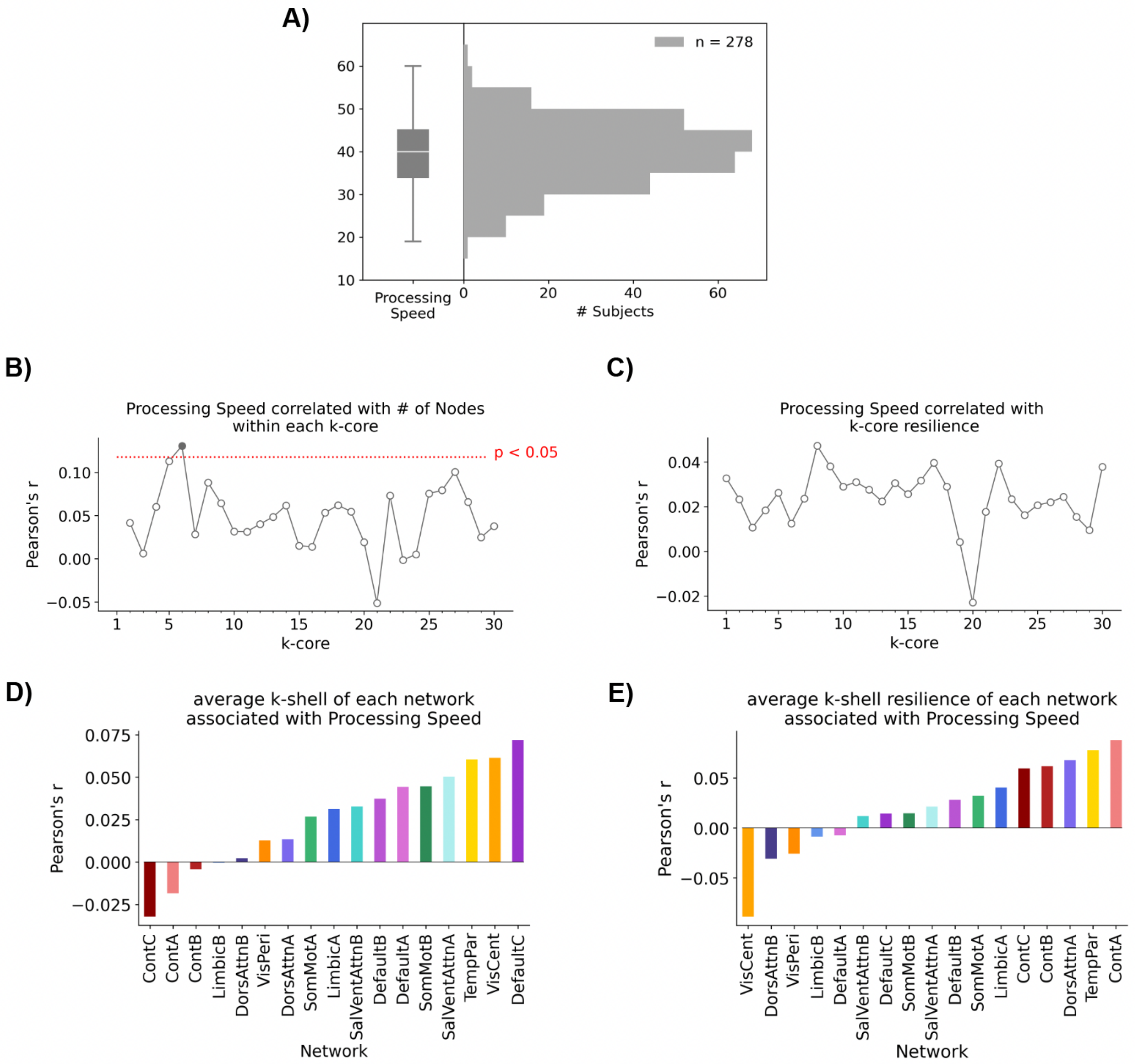


**Figure S4.** The presence, nor resilience, of robustly connected nodal communities is related to processing speed. **A)** Distributions of processing speed performance among participants. **B)** Processing speed was related to *k*-core size in only a single *k*-core (6). **C)** Processing speed was not related to resilience at any *k*-cores. **D)** Processing speed was not highly relayed to the average *k*-shell assignment for any network. **E)** Processing speed was not highly related to the *k*-shell resilience of any network.

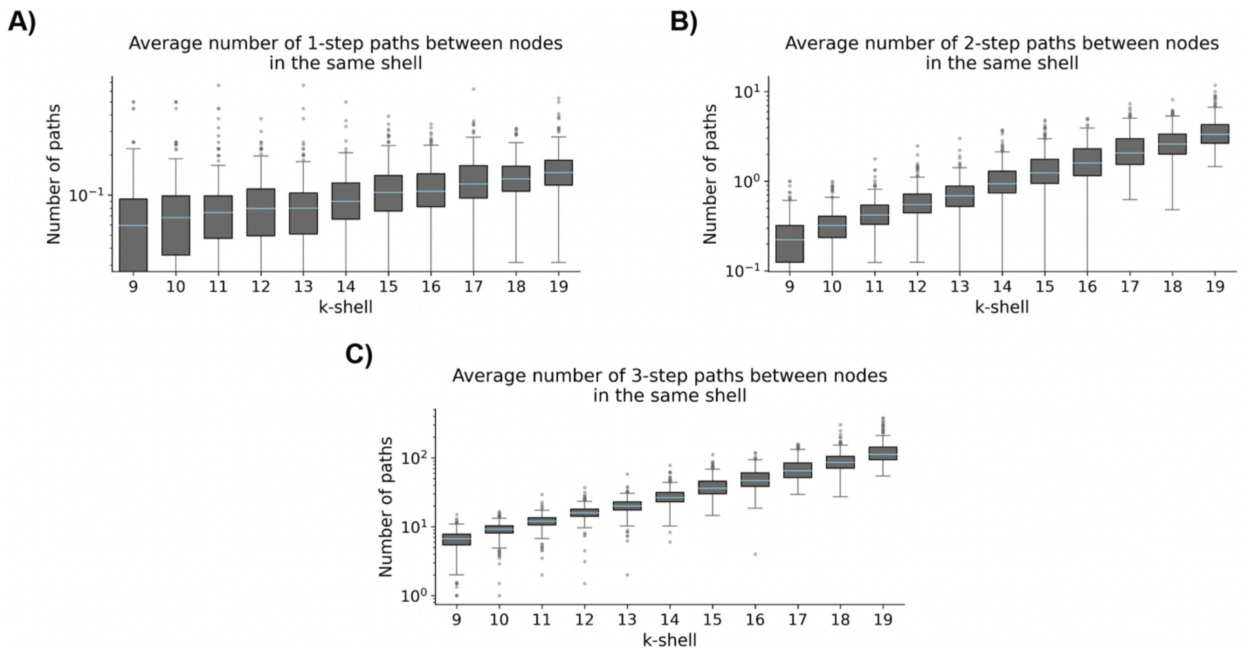


**Figure S5.** The number of redundant short paths between brain regions within the same *k*-shell increases at an approximately exponential rate as the value of *k* increases. We illustrate this for a set of *k*-shells (9-19) where every subject in our dataset had a minimum of two brain regions assigned. Each point within a distribution represents the average number of X-step paths between nodes within the respective *k*-shell for a single subject. We focus on **A)** 1-step paths (direct edges between two brain regions), **B)** 2-step paths, and **C)** 3-step paths. The y-scale in each plot has been log transformed. No attempt at quantifying these relationships were made as this was not the focus of our study, but an approximately exponential trend can be observed in each case, with this being the most apparent for 3-step paths.

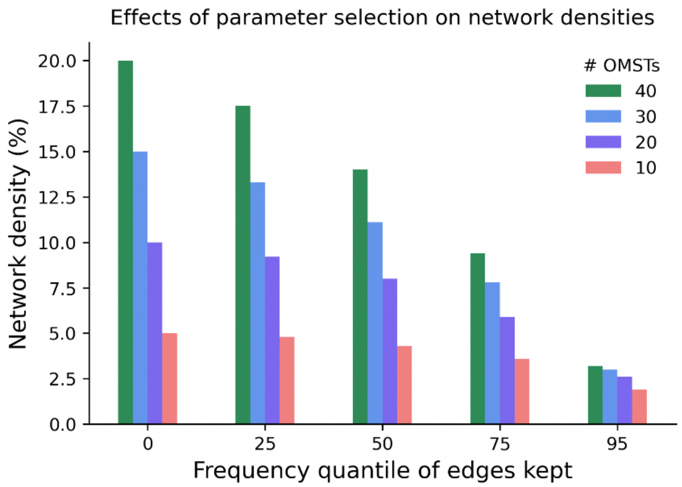


**Figure S6.** Effects of parameter selection on network densities. This can alternatively be thought about as filtering based on how much variability one would like to keep among edge pairs at the group level. For the analyses performed within the main text, we used the parameter settings of 20 OMSTs thresholded with edges thresholded if they occurred at a rate less than the 50$^{th}$ quantile of edge frequencies.

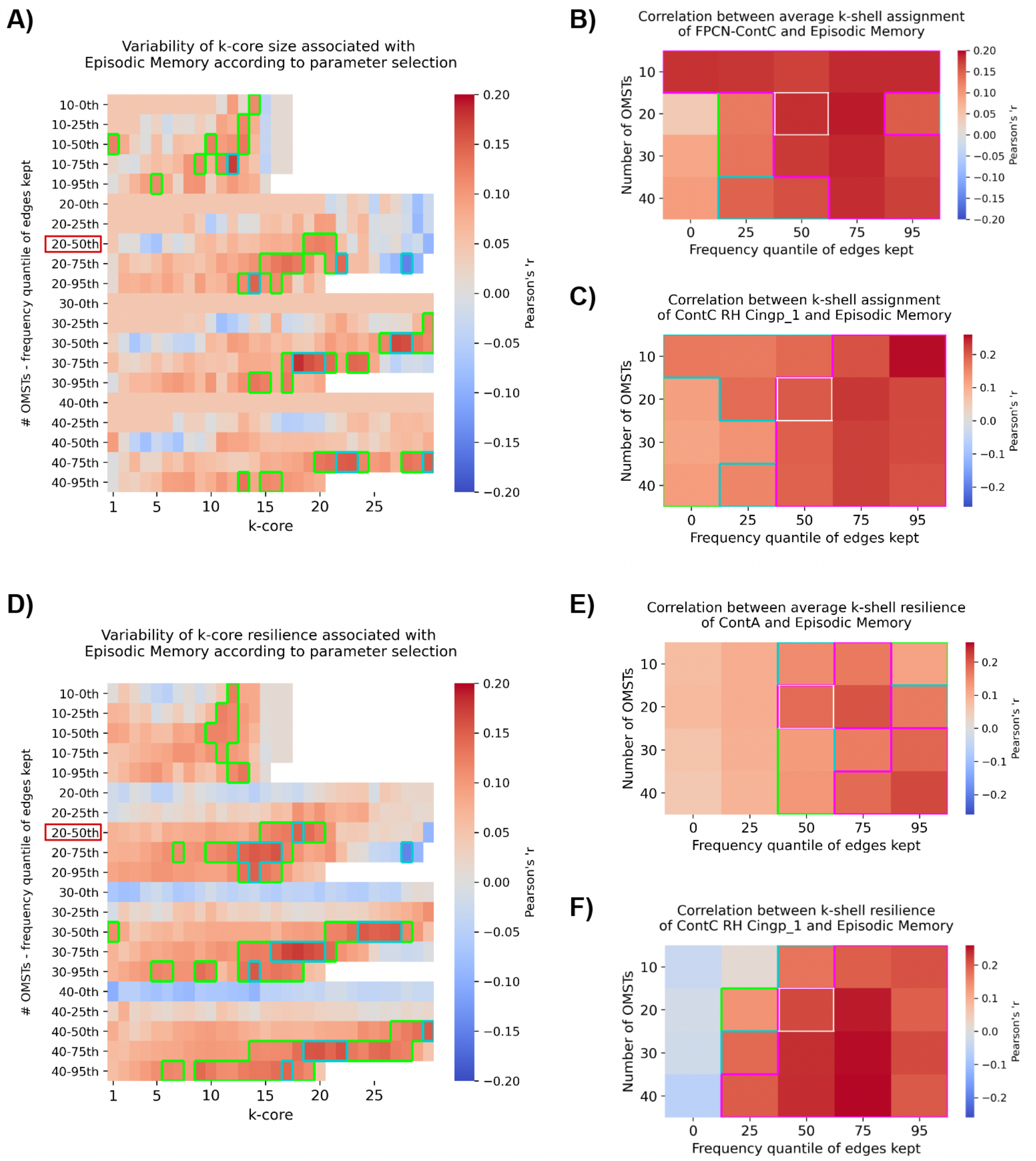


**Figure S7.** Network features associated with episodic memory are relatively insensitive to parameter selection. **A)** Variations in *k*-core size associated with episodic memory parameter selection. **B)** Variations in the association between average *k*-shell assignment of FPCN-ContC and episodic memory across parameter settings. **C)** Partial correlations between shell assignment of the ContC right hemisphere posterior cingulate region (MNI coordinates: (7, -44, 20)) and episodic memory across parameter settings. **D)** Variations in *k*-core resilience associated with episodic memory across parameter selections. **E)** Partial correlations between *k*-shell resilience of ContA and episodic memory across parameter settings. **F)** Partial correlations between *k*-shell resilience of ContC right hemisphere posterior cingulate region (MNI coordinates: (7, -44, 20)) and episodic memory across parameter selections. Areas bounded by green, and cyan lines denote significance levels of $p$-value $< 0.05$, and $p$-value $< 0.01$, respectively. Areas bounded by the magenta borders denote significance after correction for multiple comparisons. The red boxes on the left of **A** and **D** and the white boxes in **B**, **C**, **E**, and **F**, indicate the parameter settings used for the analyses in the main text. Age is included as a covariate in all analyses performed above.

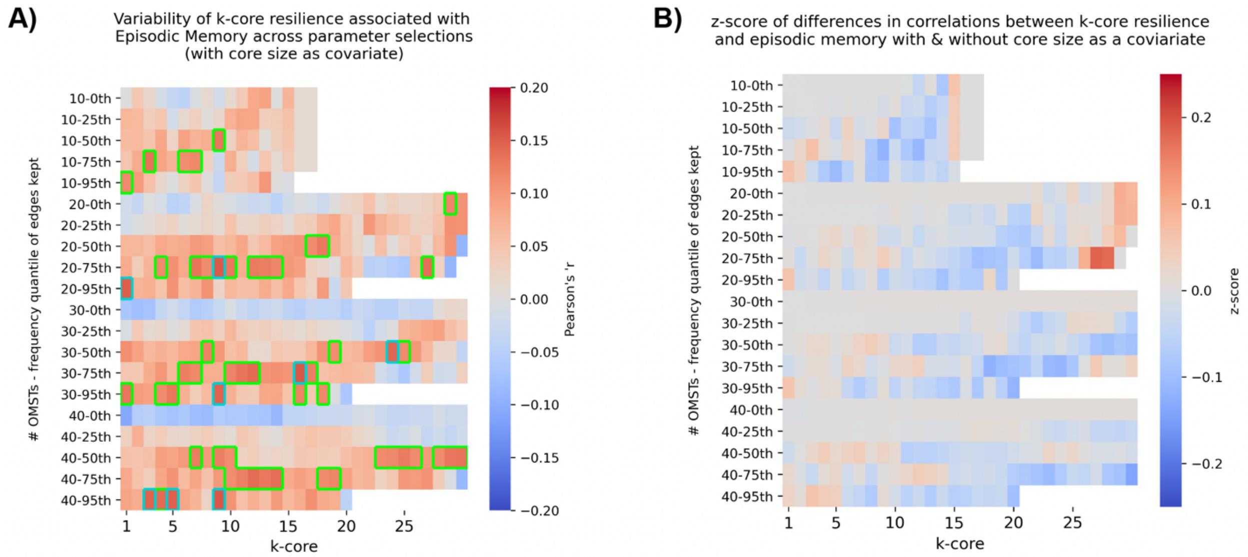


**Figure S8.** Including initial core size as a covariate when associating *k*-core resilience with episodic memory reduces the number of significant relationships observed but does not significantly change any of the correlations. **A)** Variations in *k*-core resilience associated with episodic memory with age and initial core size as covariates across parameter selections. **B)** *z*-score of the differences between *k*-core resilience and episodic memory before and after inclusion of initial core size as a covariate (with size covariate – w/o size covariate).

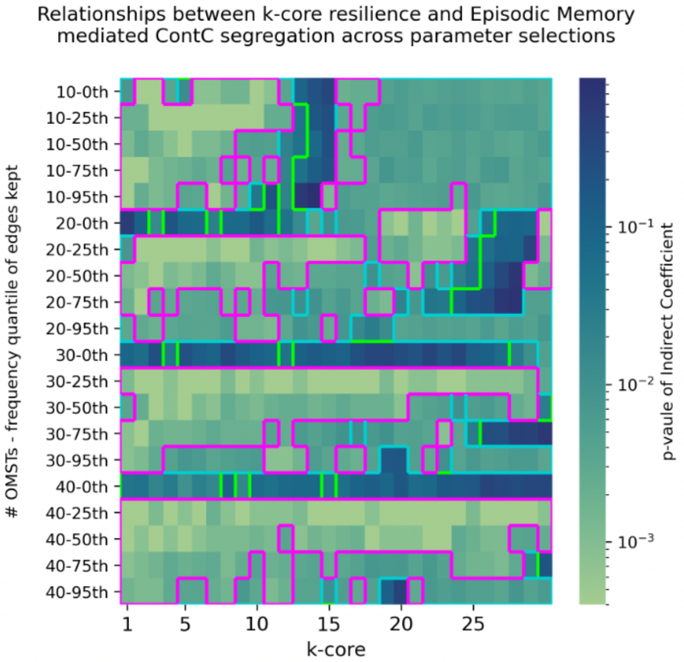


**Figure S9.** Segregation of ContC mediating relationships between resilience of early *k*-cores and episodic memory is consistent across parameter settings. The indirect coefficients of these relationships were difficult to visualize in this format. Instead, we show the significance levels of the indirect coefficients. Areas bounded by green, and cyan lines denote significance levels of p-value < 0.05, and p-value < 0.01, respectively. Areas bounded by the magenta borders denote significance after correction for multiple comparisons (corrected p-value < 0.05). Age is included as a covariate in all mediation analyses performed above. Note: results may vary slightly for the mediation analyses in main text at the 20 – 75th parameter settings as they were done with different random seeds.

| Partial correlations between Network Segregation and Episodic Memory (w/ Age covariate) | | | | |
|---|---|---|---|---|
| | Pearson's r | p-value | CI95%_L | CI95%_U |
| ContA | -0.05734 | 0.30727 | -0.17 | 0.05 |
| ContB | 0.04196 | 0.45522 | -0.07 | 0.15 |
| **ContC** | 0.17783 | 0.00143 | 0.07 | 0.28 |
| DefaultA | 0.03876 | 0.49027 | -0.07 | 0.15 |
| DefaultB | 0.06744 | 0.22968 | -0.04 | 0.18 |
| DefaultC | 0.00542 | 0.92319 | -0.1 | 0.12 |
| DorsAttnA | 0.06797 | 0.22604 | -0.04 | 0.18 |
| DorsAttnB* | 0.1108 | 0.04802 | 0 | 0.22 |
| LimbicA | 0.08234 | 0.14228 | -0.03 | 0.19 |
| LimbicB | 0.02197 | 0.69591 | -0.09 | 0.13 |
| SalVentAttnA | -0.02808 | 0.61736 | -0.14 | 0.08 |
| SalVentAttnB | 0.02285 | 0.68436 | -0.09 | 0.13 |
| SomMotA | 0.06643 | 0.23678 | -0.04 | 0.17 |
| SomMotB | 0.00567 | 0.91969 | -0.1 | 0.12 |
| TempPar | 0.03263 | 0.56148 | -0.08 | 0.14 |
| VisCent | 0.08963 | 0.11011 | -0.02 | 0.2 |
| VisPeri | 0.04978 | 0.37554 | -0.06 | 0.16 |

**Table S1.** Partial correlations between Network Segregation and Episodic Memory (with age included as a covariate) for all 17 subnetworks. ContC Segregation was significantly associated with episodic memory after correcting for multiple comparisons (corrected p-value $< 0.05$). However, segregation of DorsAttnB showed a marginal correlation with Episodic Memory ($p < 0.05$).
